# habCluster: Identifying Geographical Boundary among Intraspecific Units Using Community Detection Algorithm in R

**DOI:** 10.1101/2022.05.06.490926

**Authors:** Chengcheng Zhang, Juan Li, Biao Yang, Qiang Dai

## Abstract

Conservation management for a species generally rests on intraspecific units, while identification of their geographic boundaries is necessary for the implementation. Intraspecific units can be discriminated using population genetic methods, yet an analytical approach is still lacking for detecting their geographic boundaries. Here, based on landscape connectivity, we present a geographical boundary delineation method, habCluster, using community detection algorithm. Community detection is an algorithm in graph theory used to identify clusters of highly connected nodes within a network. We assume that the habitat raster cells with better connections tend to form a continuous habitat patch than the others, thus making the range of an intraspecific unit. The method is tested on grey wolf (*Canis lupus*) habitat in Europe and on giant panda (*Ailuropoda melanoleuca*) habitat in China. The habitat suitability for grey wolves and giant pandas were evaluated using species distribution modeling. Each cell in the habitat suitability index (HSI) raster is treated as a node and directly connected with its eight neighbor cells. The edge weight between nodes is the distance between the center of them weighted by the average of their HSI values. We implement habCluster using R programming language with inline C++ code to speed up the computing. The geographical clusters detected were compared with the HSI maps for both species and with the nature reserves for giant panda. We found the boundaries of the clusters delineated using habCluster could serve as a good indicator of habitat patches, and they match generally well with nature reserves in the giant panda case. habCluster can provide spatial analysis basis for conservation management plans such as monitoring, translocation and reintroduction, as well as for population structure research.

## 1 INTRODUCTION

We live amid a global wave of biodiversity loss, as signaled by mass species extinction, and underlying population extirpations and declines in local abundance (Dirzo et al., 2014; Johnson et al., 2017). The disappearance of populations is a prelude to species extinction (Ceballos and Ehrlich, 2002), thus the seriousness of Earth’s sixth mass extinction maybe underestimated if we overlook the declines of populations (Ceballos et al., 2017). As compare with species extinctions that are of great evolutionary importance, the declines of populations generally cause greater immediate impacts on ecosystem functions and services (Ceballos et al., 2017; Brodie et al., 2021). Local population is also the center of intraspecific conservation actions because, in practice, conservation implementations are often achieved through the protection of natural populations and their habitats. Conservation can prevent isolated and small population from entering extinction vortex due to bottlenecks and inbreeding depression (Frankham, 1998; Fagan and Holmes, 2006). However, it is usually difficult to define the boundaries of intraspecific units, despites its importance in conservation managements. Although populations and habitats are dynamic, management units must be based on certain geographic ranges to delineate entities for monitoring population and regulating the effects of human activities upon them.

Genetic methods are frequently used to delineate conservation units, for instance, the term ‘evolutionarily significant units (ESUs)’ was developed to ensure populations with unique evolutionary potential is recognized and protected, and ‘management units (MUs)’ is applied to identify functionally independent populations that connected by low levels of gene flow (Moritz, 1994). The concepts and criteria for ESUs and MUs seem logically and theoretically valid, but it remains to be seen whether they can lead to improved conservation. As the identification of ESUs and MUs is susceptible to error because of insufficient sampling: too few individuals, too few loci (Moritz, 1994), unable to detect key loci present substantial ecological and societal benefits (Prince et al., 2017), unable to account for linkage between loci or integrate data on both neutral and adaptive loci could lead to failure to recognize important genetic patterns (Allendorf et al., 2010; Funk et al., 2012). Besides, the delineation of MUs has often been misguided by focusing on rejecting panmixia rather than basing upon the amount of population genetic divergence (Palsbøll et al., 2007). Defining an ESU or MU in an operational or pragmatic sense rather than in an academic or semantic sense, however, is challenging. Because, practically, it is always necessary to define a specific range in which actions could be carried out, so conservation activities are also geographically bounded. Therefore, the delineation of a spatial boundary for population is of great significance, especially when serve as basis for defining conservation management units.

Seeking to delineate the spatial boundaries of intraspecific units from georeferenced habitat data, we developed a novel method, habCluster, which is based on the community detection algorithm in graph theory. Community detection in networks is a technique for finding groups within complex systems. A particular network may have multiple communities that nodes inside a community are densely connected. Community structure, i.e. the organization of vertices in clusters, can be detected and considered as fairly independent compartments of a graph (Fortunato, 2010). Detecting communities is of great importance in sociology, biology and computer science, which have been applied to real networks such as social networks (Bedi & Sharma, 2016), bank fraud (Sarma et al., 2020) and biological networks (Buhnerkempe et al., 2016; Tripathi et al., 2019).

habCluster introduces a new approach for delineating the boundaries of intraspecific units by (i) evaluate the connectivity between pairwise habitat cells in a landscape and (ii) aggregate cells that have strong connectivity with each other into one cluster. habCluster thus permits to identify multiple clusters from a landscape surface, which may indicate possible population boundaries. To illustrate habCluster, we used grey wolf (*Canis lupus*) habitats data in Europe and giant panda (*Ailuropoda melanoleuca*) habitat data in China as case studies, analyzed the clusters of habitats for the two species. The habitat suitability index (HSI) data used in this paper are either from open access or from published resources.

## 2 MATERIALS AND METHODS

### 2.1 Methodology

habCluster is implemented as an open-source R script with inline C++ codes. Source code, user manual and training data sets can be downloaded from: https://github.com/qiangxyz/habCluster. We hereafter detail the steps of habCluster, with Figure 1 illustrating the conceptual framework and Figures 2, 3 demonstrating its application in a case study.

**Figure 1.**
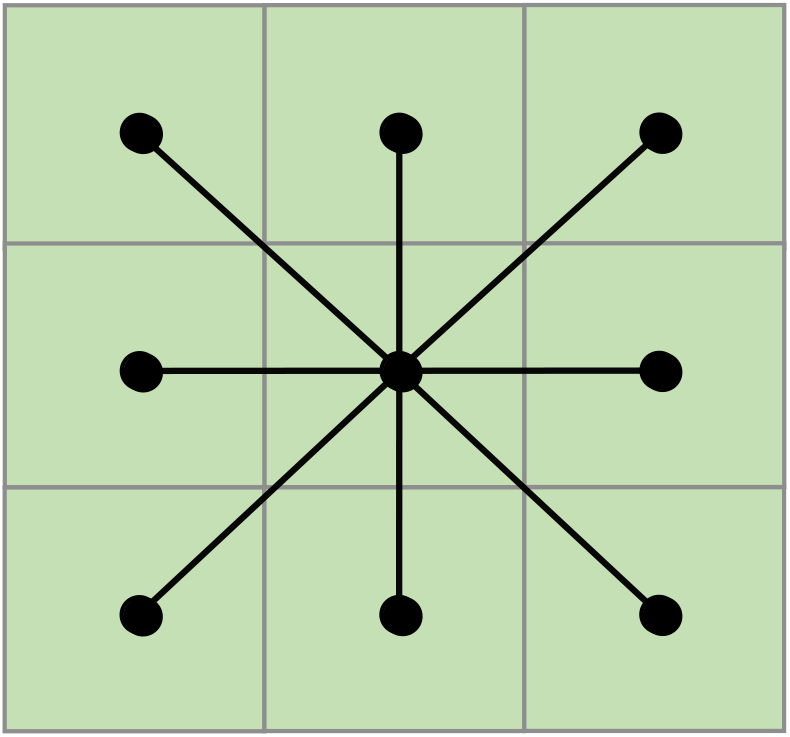
Illustration of nodes and edges in raster map. Dots represent the nodes, lines that connected dots pair represent the edges, cells in the background gird represent pixels in a raster.

**Figure 2.**
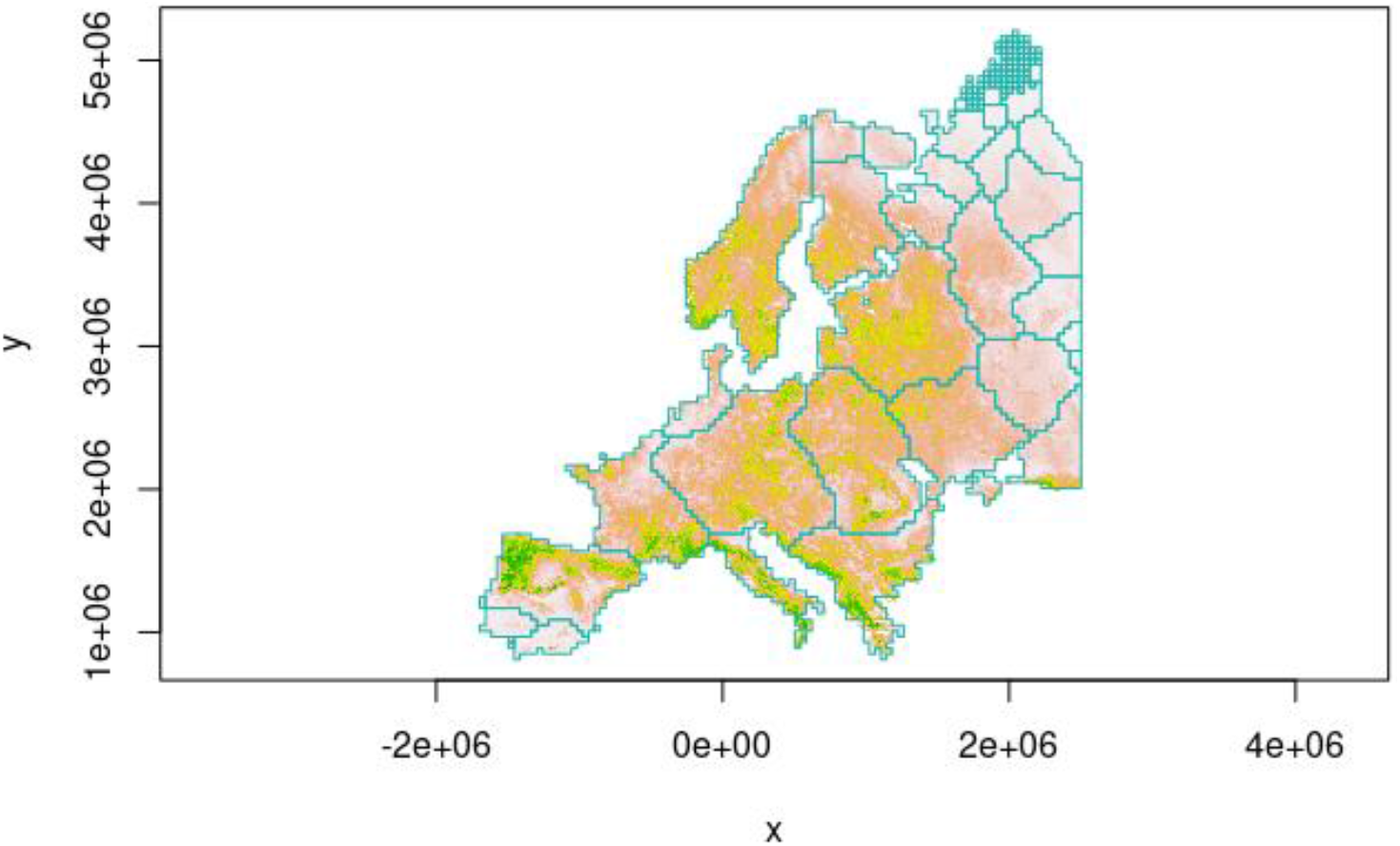
The boundaries of European grey wolf habitat clusters (light seagreen polygons) delineated from community detection is shown against the habitat suitability index map (colors from grey to green indicating from less to more suitable habitats).

**Figure 3.**
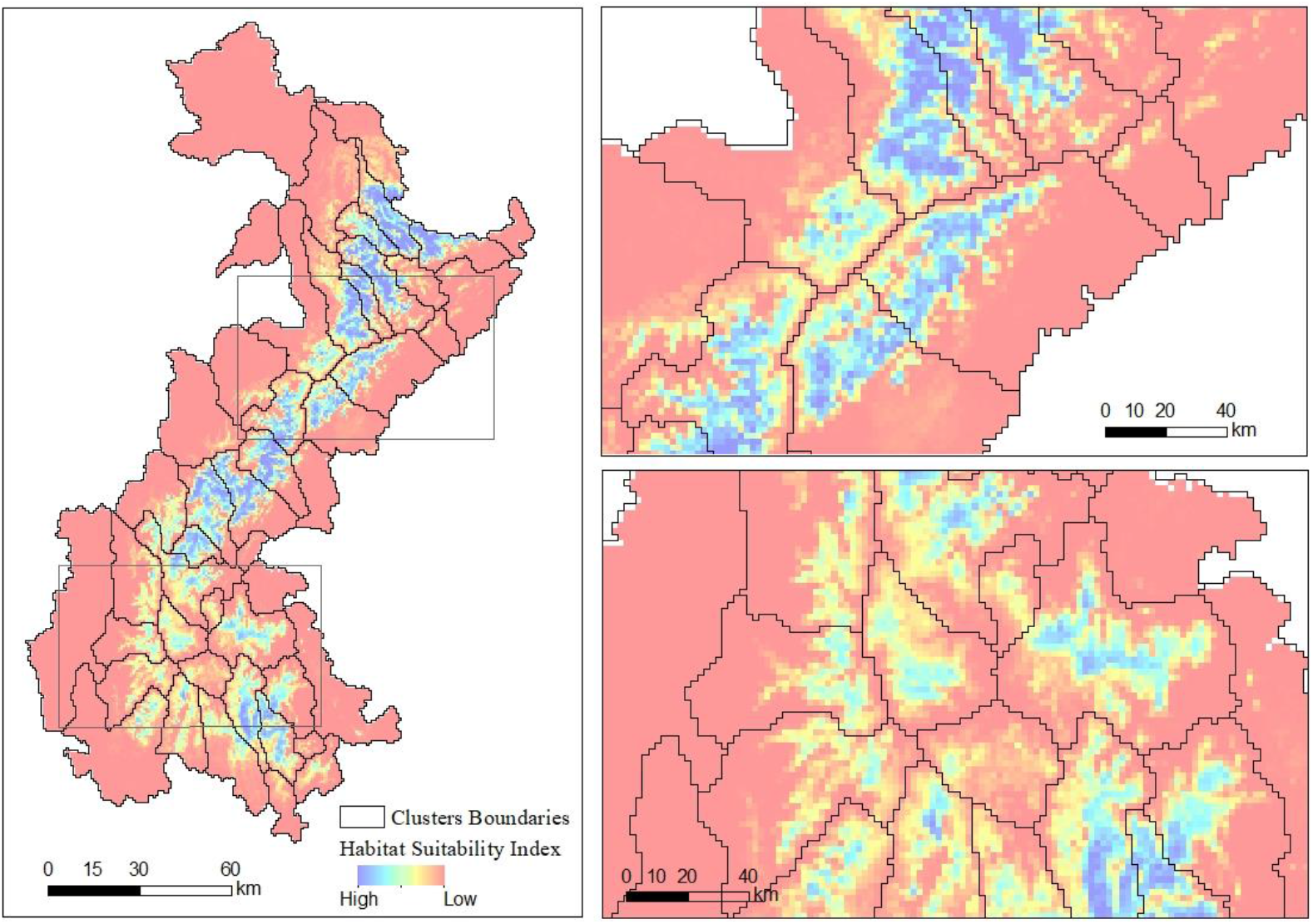
The boundaries of giant panda habitat clusters (black polygons) delineated from community detection is shown against the habitat suitability index map (colors from blue to red indicating from less to more suitable habitats).

Isolation by distance (IBD) model considers that the dispersal distance of individual is limited, so the ones are geographically close tend to be genetically more similar than the individuals that are far apart, also tend to be aggregated into a local population (McRae, 2006; Meirmans, 2012). IBD model has assumed spatial homogeneity in species’ distributions and migration rates, however, it failed to account for landscape heterogeneity, while isolation by resistance (IBR) model filled this gap (McRae, 2006). Landscape resistance, an indication of how well a landscape can be traversed by a given species, usually estimated simply as the inverse of habitat suitability (Rudnick et al., 2012). Where resistance is low (higher structural connectivity), individuals are more likely to move through the landscape, thus it is easier for the individuals to form a functionally connected population. This implies that we can divide landscape into various clusters according to its spatial connectivity, thereby the boundaries of intraspecific units can be delineated.

#### 2.1.1 Data input

The method requires a raster map which refers to how much the landscape facilitates individual movements. Our analysis requires evaluation of how strong two cells are connected, therefore, the value of habitat suitability of a cell was described as ‘smoothness’ to indicated how easy an individual can move through. Habitat suitability raster layer can be evaluated using various algorithms like MaxEnt, General Liner Model (GLM), random forest, etc. (Bai et al., 2018; Zacarias and Loyola, 2018).

#### 2.1.2 Connectivity between cells

Each cell of a HSI raster is treated as a node and directly connected with its eight neighbor cells (Figure 1), its value represents how easy to move through the cell for a species. The connection between two adjacent cells is treated as edge. The edge weight between nodes is the distance between the center of them weighted by the average of their HSI values. It indicates a strong connectivity between the two cells when they are suitable habitat, and individuals can move smoothly between them, and vice versa. Non-adjacent cells can be indirectly connected via intermediate cells, thus all cells in the entire map can be connected together.

#### 2.1.3 Calculating processes

Nodes and edges are created and added to a graph. The nodes can be clustered according to the connectivity of the edges by community detection algorithm. Those nodes that are densely connected to each other would be preferentially aggregated together, corresponding to spatially closely connected areas. qs (Eddelbuettel and Francois, 2011) to improve computing speed. We use the R package ‘igraph’ to implement community detection algorithm (Csardi and Nepusz, 2006).

#### 2.1.4 Data output

igraph outputs the community identification numbers that all nodes belong to. The information is stored as a raster file, so the boundaries of clusters could be delineated.

### 2.2 Case studies

To illustrate the uses of habCluster, we analyzed the habitat clusters from two biological datasets.

#### 2.2.1 Grey wolf dataset

Distribution points of European wolves were obtained from open dataset (GBIF Secretariat, 2022) and roughly adjusted with published data (Stronen et al., 2013). Bioclimatic factors (Fick and Hijmans, 2017), elevation (Jarvis et al., 2008), and land cover (ESA, 2017) were used as environmental variables to compute the wolf HSI map using MaxEnt.

Install the development version of habCluster from GitHub.

~~~
devtools::install_github(“qiangxyz/habCluster”)
~~~

Load the packages (Edzer Pebesma, 2018; Hadley Wickham et al., 2021; Hijmans Robert J., 2021) needed for analysis.

~~~
library(sf)
library(raster)
library(dplyr)
library(habCluster)
~~~

Load the HSI data of European wolf. The HSI values in the raster cells indicate how smoothly the wolves can move in the cells, and can be used to represent the habitat connection between cells.

~~~
hsi.file = system.file(“extdata”,”wolf3_int.tif”,package=“habCluster”)
wolf = raster(hsi.file)
~~~

Find habitat clusters using Leiden algorithm. Raster for habitat suitability was resampled to a resolution of 40 by 40 km (40000 m) to reduce calculation amount. Set cluster_resolution_parameter to 0.02 to control the cluster size. Note that the parameter of `cellsize` controls the spatial scale analysis is performed, while the parameter of `rp` is used to control cluster size.

~~~
clst = cluster(wolf, method = cluster_leiden, cellsize = 40000, rp = 0.02, silent = FALSE)
~~~

Plot clusters result with European wolf HSI raster as base map.

~~~
image(wolf, col = terrain.colors(100, rev = T), asp = 1)
boundary = clst$boundary
plot(boundary$geometry, add=TRUE, asp=1, border = “lightseagreen”)
~~~

Or discard small patches before plotting.

~~~
image(wolf, col = terrain.colors(100, rev = T), asp = 1)
boundary$area = st_area(boundary)%>%as.numeric
boundary = boundary %>% filter(area > 40000*40000)
plot(boundary$geometry, add=TRUE, asp=1, border = “lightseagreen”)
~~~

#### 2.2.2 Giant panda dataset

The giant panda presence records were obtained from surveys covering an area of 330,000 km^2^ and 44 giant panda reserves. The HSI map were evaluated using MaxEnt modeling, with 1421 valid species presence sites and 29 spatially explicit environmental variables (Qing et al., 2016). The example data is a raster resampled from the giant panda HSI map, whose values range from 0 to 1.

Install the development version of habCluster from GitHub.

~~~
devtools::install_github(“qiangxyz/habCluster”)
~~~

Load the HSI data of giant panda.

~~~
panda = read_stars(“./hsi_panda.tif”)
~~~

Louvain clustering algorithm is used to detect clusters. Habitat suitability raster was resampled to a resolution of 2 km by 2 km (2000 m). Note that the parameter of `cellsize` controls the spatial scale analysis is performed, while the parameter of `rp` is used to control cluster size.

~~~
clst = cluster(panda, method = cluster_louvain, cellsize = 2000, rp = 0.02, silent = FALSE)
~~~

Plot clusters result with giant panda HSI raster as base map.

~~~
image(panda, col = terrain.colors(100, rev = T), asp = 1)
boundary = clst$boundary
plot(boundary$geometry, add=TRUE, asp=1, border = “black”)
~~~

Finally, the resulting habitat clusters were overlapped with the boundaries of giant panda nature reserves.

## 3 RESULTS

### 3.1 Grey wolf dataset

The boundaries of European grey wolf habitat clusters delineated from community detection is shown in Figure 2.

### 3.2 Giant panda dataset

The boundaries of giant panda habitat clusters delineated from community detection is shown in Figure 3. These boundaries divisions show well performance in detecting areas that are separated by low HSI values, thus could serve as a good indicator of habitat patches and intraspecific units.

The majority of giant panda nature reserves share similar spatial boundaries with the delineations from our community detection results, although a few exceptions do exist (Figure 4).

**Figure 4.**
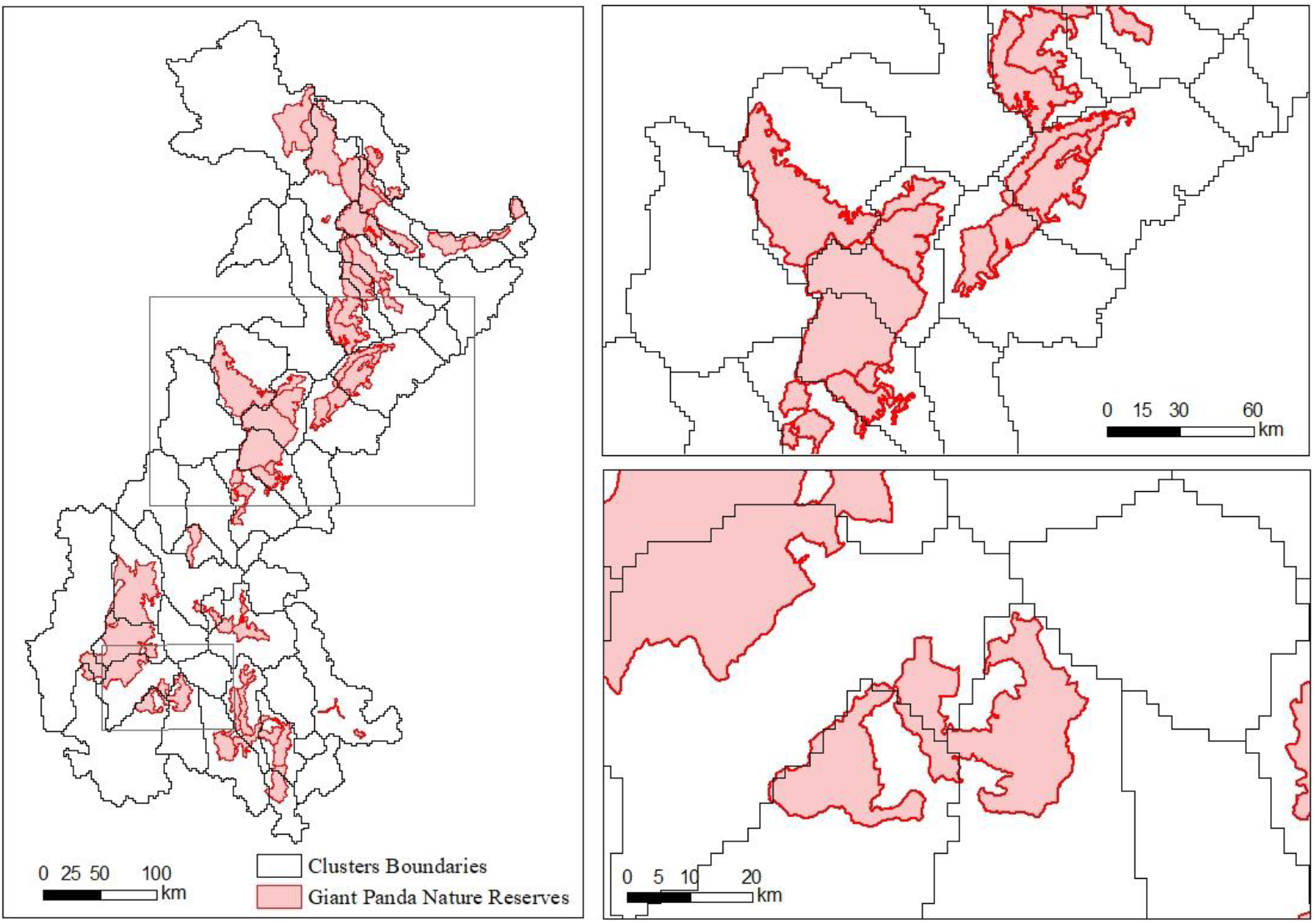
The boundaries of giant panda habitat clusters (black polygons) delineated from community detection and the nature reserves (red polygons).

## Discussion

We presented a new method to delineate the boundaries of intraspecific units. habCluster provides great potentials in conservation management as it gives pragmatic use in defining conservation units on certain geographic ranges. As measuring habitat connectivity and suitability has increasingly become a routine objective of researchers and policy makers (Rudnick et al., 2012), one step further to population boundaries delineation is promising. The boundaries of clusters detected with habCluster could serve as a good indicator of habitat patches, giving convincing divisions on intraspecific units such as population, subpopulations, or local populations, and corresponding entities for regular monitoring.

habCluster provides a spatial analysis tool for population ecology research, as well as for delineation of conservation management units and realization of effective managements. Specifically, it can be used in planning or improving protected areas and conservation strategies as it reveals the current status of habitat fragmentation. Our result of European wolf habitat clusters shares similarity with the population clusters derived from genetic study (Stronen et al., 2013). Which shows great potential in applying habCluster for conservation management. For planning, habCluster can provide insights in where conserved areas should be established so actions could be taken accordingly. For improving protected area network, it can be used to evaluate the conservation effectiveness from the perspectives of threats and outcomes. As shown in our giant panda case, the overlaps between clusters boundaries and nature reserves suggested some nature reserves cover multiple clusters (Figure 4. bottom-right). In this case, landscape corridors could be designed to maintain or improve the connectivity between local populations.

An additional caveat is that scales matter when applying habCluster for conservation planning or assessment. In choosing the resolutions of HSI map used for community detection, the behavioral and ecological characteristics of the target species should be taken into considerations, so do the requirements of conservation managements, thus the scale of analysis with practical significance can be determined.

## Data availability

The codes and example data for this study are available on GitHub: https://github.com/qiangxyz/habCluster.

## Author Contributions

QD, and BY designed the experiments. QD, CZ and JL performed the experiments. CZ, JL, BY and QD analysed the results and wrote the manuscript.

## Acknowledgments

We would like to thank the team of the Fourth National Giant Panda Survey (Sichuan) for providing us the case study data used in this work.

